# Tension-induced adhesion mode switching: the interplay between focal adhesions and clathrin-containing adhesion complexes

**DOI:** 10.1101/2024.02.07.579324

**Authors:** Umida Djakbarova, Yasaman Madraki, Emily T. Chan, Tianyao Wu, Valeria Atreaga-Muniz, A. Ata Akatay, Comert Kural

**Author notes:** Present address: National Metrology Institute, Scientific and Technical Research Council of Turkey, 41470 Gebze-Kocaeli, Turkey (A. Ata Akatay).

## Abstract

Integrin-based adhesion complexes are crucial in various cellular processes, including proliferation, differentiation, and motility. While the dynamics of canonical focal adhesion complexes (FAs) have been extensively studied, the regulation and physiological implications of the recently identified clathrin-containing adhesion complexes (CCACs) are still not well understood. In this study, we investigated the spatiotemporal mechanoregulations of FAs and CCACs in a breast cancer model. Employing single-molecule force spectroscopy coupled with live-cell fluorescence microscopy, we discovered that FAs and CCACs are mutually exclusive and inversely regulated complexes. This regulation is orchestrated through the modulation of plasma membrane tension, in combination with distinct modes of actomyosin contractility that can either synergize with or counteract this modulation. Our findings indicate that increased membrane tension promotes the association of CCACs at integrin αVβ5 adhesion sites, leading to decreased cancer cell proliferation, spreading, and migration. Conversely, lower membrane tension promotes the formation of FAs, which correlates with the softer membranes observed in cancer cells, thus potentially facilitating cancer progression. Our research provides novel insights into the biomechanical regulation of CCACs and FAs, revealing their critical and contrasting roles in modulating cancer cell progression.

## Introduction

Cellular adhesion to the extracellular matrix (ECM) is a complex, tightly regulated process essential to most cellular functions, such as proliferation, differentiation, and motility(1–3). This adhesion is primarily mediated by membrane bound integrin receptors, which assemble into highly organized structures known as adhesion complexes. These structures integrate signals from both the internal and external environments of the cell, serving as pivotal points for signal transduction (4–6). Although cells exhibit a variety of integrin-based adhesion complexes with distinct properties and roles (5–11), the overall interaction between these complexes and their collective influence on cellular behaviors is not fully understood. Investigating the spatiotemporal regulations of adhesion complexes, especially by mechanical cues at the plasma membrane, is crucial for comprehensive understanding of their roles in processes involving extensive membrane remodeling, such as cell proliferation, migration, and differentiation.

Focal adhesions (FAs) are the most extensively studied integrin-based adhesion complexes that link the ECM and cytoskeleton, playing a critical role in mechanotransduction and cellular dynamics. The dynamic regulation of FAs is influenced by both the chemical and physical properties of the ECM, as well as the contractile forces from actomyosin networks (7,12–17). Different actin structures within cells lead to various contractile behaviors and are responsible for distinct cellular adhesions. Aligned stress fibers, higher-order cytoskeletal structures made of cross-linked actin filaments and myosin motor proteins, induce dipole-like contractility (18,19). This contractility is key in the assembly and stabilization of FAs. In contrast, cortical actins, which are disordered actomyosin networks beneath the plasma membrane, promote isotropic contraction and support cell-cell adhesion (20–22). Shifts in actin organization from stress fibers to cortical structures lead to the dissociation of FAs, highlighting the critical balance of actomyosin organization and modes of contractility in FA regulation (23–25).

Recent studies revealed a novel class of integrin-based adhesion complexes that are enriched in clathrin (26–30). In the canonical scheme, clathrin, along with its adaptor proteins, creates curved and dynamic clathrin-coated pits to drive the internalization of membrane components, including FAs, through a process known as clathrin-mediated endocytosis (CME) (31–35). However, not all clathrin-coated structures (CCS) mature into pits and invaginate; some remain flat and reside on the membrane for long periods of time (36–40). These remarkably static and larger arrays of CCS are often termed as clathrin-coated plaques, flat clathrin lattices or clathrin sheets and exclusively localized on the ventral/adherent surface of cultured cell (29,41). Besides their endocytic roles, they are known to have pleiotropic non endocytic functions. For instance, they are observed in close proximity to the substrate and directly associate with integrins and ECM substrates (28,42–49), indicating their putative role in cell adhesion. Furthermore, they function as mechanotransduction units by sensing the extracellular milieu and facilitating signaling via clustering plasma membrane receptors, therefore regulating distinct tissue specific cellular processes (45,50–52). Given their adhesive and mechanosensitive properties, we will refer to these structures as clathrin-containing adhesion complexes (CCACs) throughout the remainder of this article.

Although CCACs share many features with FAs as adhesive entities, they lack classical components of adhesion complexes and as well as a link to the actomyosin cytoskeleton(53). Thus, CCACs are not directly influenced by actomyosin contractility; instead, they are affected by the biomechanical properties of the ECM and the local membrane tension. Therefore, impediments of endocytosis due to physical or structural constraints including the size of cargo molecules(54–56), substrate rigidity (Baschieri et al., 2018), strength of cell adhesion sites (58,59) and failure to overcome local forces to generate curvature due to high membrane tension prolong the presence of CCS on the plasma membrane and promote CCAC formation (60–64). Restoring the plasma membrane tension facilitates formation of clathrin pits and triggers CCAC dissociation (60–64), indicating membrane tension acts as a potent and reversible regulator of CCAC formation and stability. Taken together, our understanding of cellular adhesion reveals a complex landscape where two distinct mechanical forces—plasma membrane tension and stress fiber-based contractile tension—uniquely regulate the formation and stability of CCACs and FAs, respectively.

Despite their distinct composition and dynamics, many adhesion complexes are functionally and spatially linked structures (65–70). Perturbation in one structure often promotes the formation of the other(s) through redistribution of integrin receptors (65,71–74), indicating complementary utilization of various adhesion complexes. In the same way, depleting FAs through changes in actomyosin contractility or reducing the expression of FA components increases CCAC formation and colocalization with integrin αVβ5, while disruption of CCACs promotes an increase in size and number of FAs and their integrin αVβ5 content. This competitive conditional existence is most evident during mitosis, where CCACs predominate in the absence of FAs (75,76). The absence of FAs during the M phase can be attributed to loss of stress fiber-based contractility due to the reorganization of actins into a cortical arrangement(77–79). This cortical actin, tethered to the plasma membrane by membrane-to-cortex adhesion complexes, underlies the heightened membrane tension, which in turn, triggers the formation of CCACs (80–86).

This suggests that crosstalk between actin network organization and plasma membrane tension dictates the context-specific and complementary employment of CCACs and FAs. For comprehensive understanding of reciprocal regulation between cell adhesion and cell cycle progression, it is crucial to establish whether the regulatory mechanisms observed during mitosis persist throughout the cell cycle, potentially influencing phase specific utilization of adhesion complexes. Moreover, while the relationship between membrane tension and actomyosin contractility is well-documented in the context of cell spreading and migration(87–90), the precise coordination of adhesion complexes by different mechanical forces during these processes require further investigation.

In this study, we explain the spatiotemporal regulation of adhesion complexes by fluctuations in mechanical forces during the cell cycle and their impact on proliferation, spreading and migration. Our findings reveal that FAs and CCACs are mutually exclusive, their expression being intricately regulated by cell cycle dependent modulation of membrane tension and transitions of actomyosin contractility modes. Significantly, our data emphasize the competitive dynamics between CCACs and FAs for integrin αVβ5 adhesion sites: elevated membrane tension and cortical actin arrangement predisposes CCACs towards αVβ5, while reduced tension and stress fiber formation facilitates αVβ5 association with FAs. This competitive binding to αVβ5 adhesion sites critically governs oncogenic cellular behaviors, including cancer cell proliferation, spreading, and migration. Overall, our study provides novel insights into the biomechanical regulation of CCACs and FAs, elucidating their respective roles as suppressors and promoters of cancer progression.

## RESULTS

### 1. CCAC and FA are inversely regulated throughout the cell cycle

To observe the regulation of CCAC and FA structures during cell cycle, we transfected genome-edited eGFP tagged AP-2 expressing SUM159 triple-negative breast cancer cells with mCardinal tagged paxillin. Cells were synchronized to M phase, harvested with the mitotic shake-off method and imaged with spinning disc confocal microscopy during cell cycle progression. Our results supported the notion that these structures were interchangeably employed; the expression of AP-2 marked CCACs and paxillin marked FAs swung inversely during the entire cell cycle (Figure 1a). AP-2 GFP labels all CCS including CCACs, therefore it is critical to establish a reliable method for distinguishing CCACs. Given their longevity and larger size, CCACs were quantified using specific thresholds applied to both the intensity (>150 a.u.) and lifetime (>120 seconds) of the tracked CCS particles (Figure S1a). We validated the reliability of our discrimination method by comparing the Z positions of CCS structures and found that the particles above the thresholds are indeed located closer to the substrate (Figure S1b), which is the prominent feature of CCACs(91).

**Figure 1:**
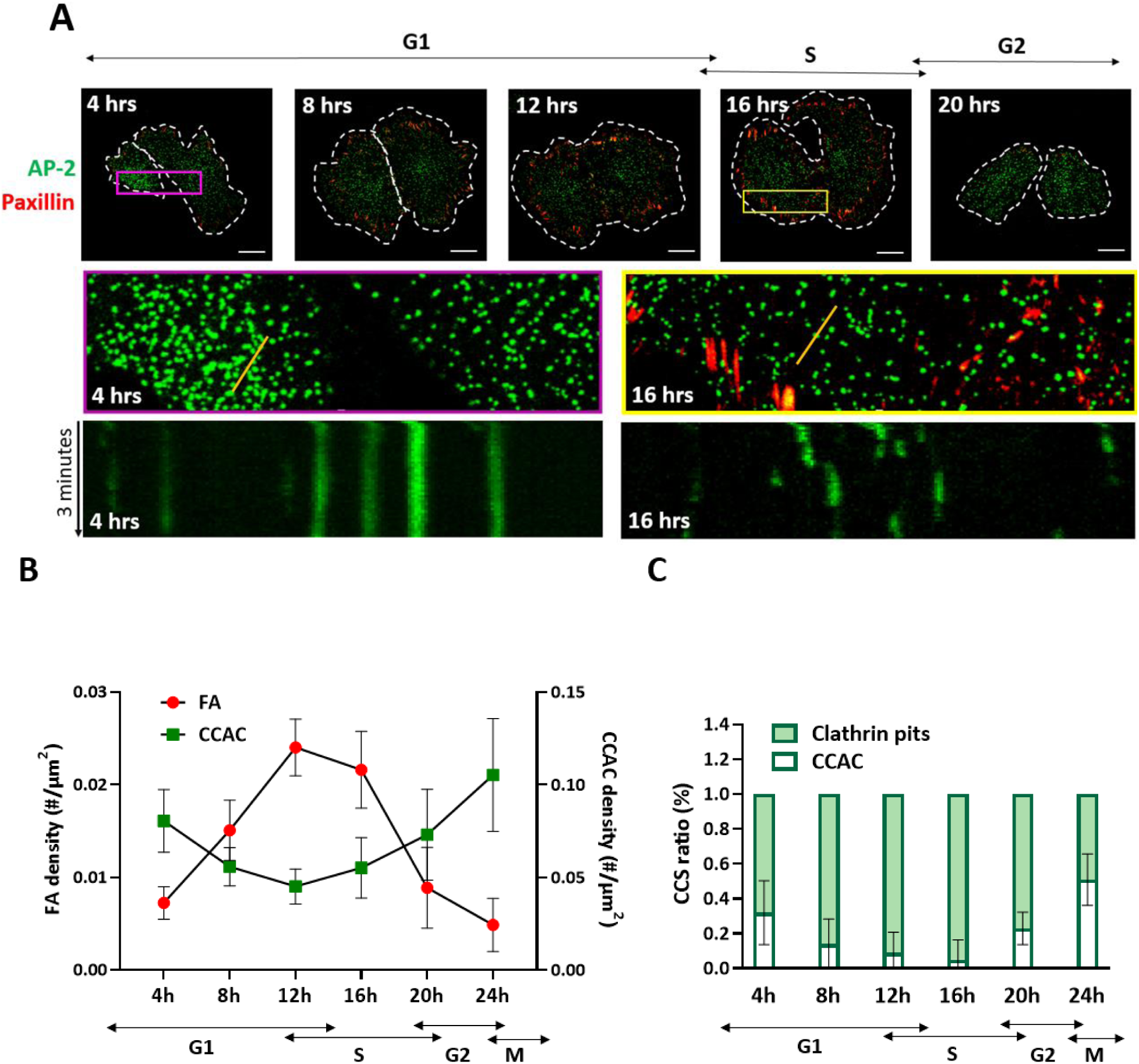
CCACs and FAs are inversely regulated during Interphase. **A**) Representative images of two daughter cells expressing AP-2 labeled CCACs (green) and Paxillin labeled FAs (red), synchronized at the M phase and imaged at various time points post-release from mitotic shake-off, corresponding to different cell cycle stages. The upper panel includes dashed lines outlining cell boundaries; scale bar: 10µm. The middle panel shows zoomed-in images of colored insets from cells in G1 (purple, 4 hrs) and S phases (yellow, 16 hrs). The lower panel presents kymographs of AP-2 labeled CCS from a representative cell during G1 (4 hrs) and S phases (16 hrs). Kymographs, derived along the regions marked with a yellow solid line in the middle panel, illustrate CME dynamics. Short streaks indicate rapid endocytic events, with streak length increasing as endocytosis slows. **B**) Line plot depicting the density of FAs and CCACs across different cell cycle stages, including standard deviation (SD). **C**) Bar graph representing the average percentage (±SD) of clathrin-coated pits or CCACs at various cell cycle time points, based on data from three or more independent experiments.

Once we optimized the quantification approaches, we compared the abundance of FAs and CCACs during cell cycle stages. We found out that CCAC expression was not limited to the M phase; instead, it slowly decreased towards the S phase and increased back before the G2/M phase (Figure 1b). Conversely, FAs displayed a gradual increase starting from the G1 phase, reaching a peak during the S phase, followed by a significant reduction at the G2/M phase, and ultimately becoming undetectable during the M phase (Figure 1b).

Interestingly, the level of clathrin pits demonstrated a similar profile to FAs throughout the cell cycle (Figure 1c). These findings were consistent with the observed changes in CME dynamics quantified by the standard deviation of growth rate, and initiation and conclusion rates of CCS formation (Figure S1c,d). The initiation and conclusion rates were low during early and late interphase but significantly higher during mid-interphase, indicating heightened CME dynamics at this stage.

Furthermore, we found that although the CCAC density in individually monitored cells exhibit a comparable trend throughout the cell cycle, the timing of decline and increase in CCAC density was not perfectly synchronized (Figure s1e), likely due to slight variations in the progression rate of the cell cycle. Interestingly, we also noticed that the inverse regulation of CCAC and FA expression was evident in the asynchronized cell population (Figure s1f), where transient overexpression of FAs coincided with lower expression of CCACs, and vice versa. These results further confirm that that CCAC and FAs are mutually exclusive structures, mutually inhibiting each other’s presence.

FAs significantly vary in size and dynamics (92). In addition to FA density, we found out that the morphometry (mean particle intensity and area coverage (Figure S1g) of FAs oscillates during interphase, indicating both nascent and mature focal complexes were cell cycle regulated.

In contrast to previous reports (75), we did not observe any significant changes in cellular localizations of both CCACs and FAs during interphase. To track the CCAC subcellular localization, we generated a temporal color map of CCS particles (Figure S1j). Further application of the intensity threshold and imaging the lowest possible z-position, left us with CCACs alone, by excluding smaller and dynamic pits. Although FAs were located at the cellular edges in some cells, the subcellular localizations of both CCACs and FAs during interphase were different for each cell, lacking any remarkable cell cycle-specific spatial imprint. Population of cells at particular cell cycle stages was verified by quantifying PI-stained nuclei with Flow Cytometry (Figure S1k).

Furthermore, to validate the functional role of CCACs as adhesion units and investigate their localization with integrins, we conducted immunostaining experiments. Previous research identified that integrins αVβ5, β3 and β1 reside in both FAs and CCACs (30,53,71,93–98). Our findings confirm that αVβ5 integrin exhibited colocalization with bright and large CCS located in proximity to surface, the features associated with CCACs, as well as with the FAs. However, no significant colocalizations of CCS and β3 and β1 integrins were observed (Figure S2a,b).

Lastly, it has been reported that CCAC colocalizes with αVβ5 exclusively during mitosis(76). However, we observed that αVβ5 to CCAC colocalization persists throughout the interphase in synchronized cells (Figure S2c). The degree of colocalization corresponded with the cell cycle-regulated oscillation of FA and CCAC densities (Figure S2d).

### 2. Cyclin D1-Cdk4/6 inhibition prevents CCAC to FA switch

The decrease in CCAC abundance and the onset of FA expression during the G1 phase (Figure 1b) temporally associated with the peak expression of Cyclin D1, which marks the Cyclin D1-Cdk4/6 activity (Figure S3a). Cdk4/6 has been identified as an upstream kinase in integrin signaling and is suggested to regulate integrin-based adhesion complexes(99). Moreover, cytoplasmic Cyclin D1-Cdk4 phosphorylates membrane-associated paxillin, promoting Rac1 activation and triggering cell spreading and migration(100). This evidence prompted us to hypothesize a potential role for Cyclin D1-Cdk4/6 in the adhesion mode switch from CCACs to FAs.

To investigate this hypothesis, we treated cells with PD332991, a selective Cyclin D-Cdk4/6 inhibitor(101). Our findings revealed a significant increase in CCAC density upon functional depletion of Cdk4/6, while concurrently reducing the morphometric parameters of FAs including density Figure 2a,b), intensity, and area coverage (Figure S3b). Additionally, there was a notable shift in the proportion of CCACs and clathrin pits in Cdk4/6-inhibited cells Figure 2c). Furthermore, the colocalization patterns of FA and CCAC with integrin αVβ5 also demonstrated alterations in response to functional depletion of Cdk4/6 (Figure S3c).

**Figure 2:**
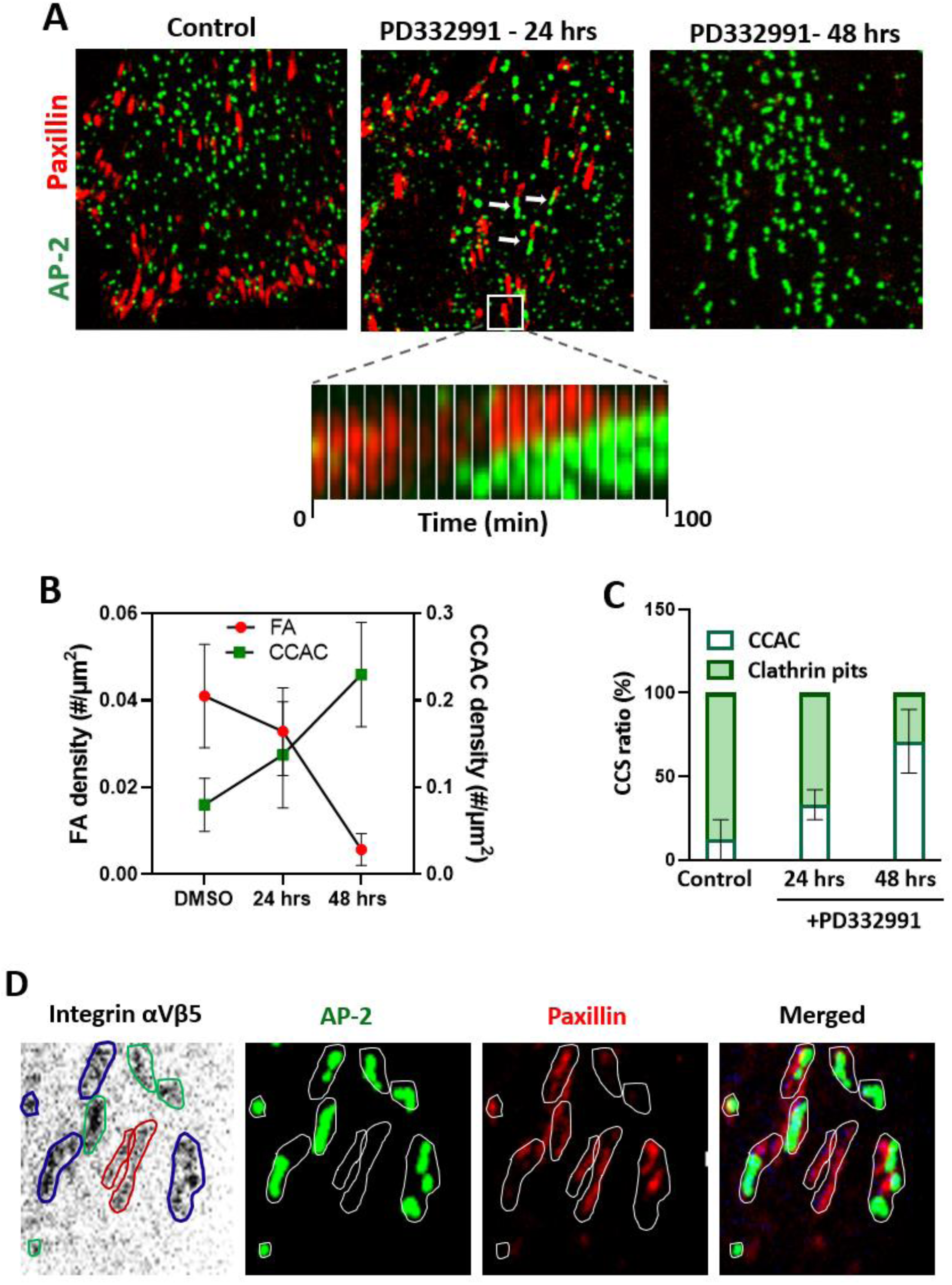
Cyclin D-Cdk4/6 is essential for CCAC to FA transition. **A**) Immunofluorescent images of cells untreated (control) and treated with PD332991 at various time points. The bottom middle panel features a montage of cropped images taken at 10-minute intervals. In the middle panel (24 hours, PD332991), white arrows highlight adhesion sites co-occupied by both CCACs and FAs. **B**) Line plot illustrating the density of FAs and CCACs in untreated (control) versus PD332991-treated cells, including standard deviation (SD). **C**) Bar graph depicting the average percentage (±SD) of clathrin-coated pits or CCACs in control versus PD332991-treated cells. Data are presented as means from three or more independent experiments. **D**) Immunostained cells, genetically edited to express AP2-GFP and stably expressing mCardinal-Paxillin, with integrin αVβ5. The integrin αVβ5 staining is inverted to enhance contrast. Blue lines indicate αVβ5 adhesion sites co-occupied by both CCAC and FA, red lines mark FA-exclusive sites, and green lines show sites occupied solely by CCAC.

Interestingly, in Cdk4/6 inhibitor treated cells, in addition to individually organized CCAC and FA structures, we identified aligned arrays of CCAC and FA presumably sharing the same adhesion sites Figure 2a, marked by arrows in the middle panel). These adhesion sites, whether exclusively occupied by individual complexes or jointly by both CCACs and FAs, exhibited linear and radially asymmetric configurations. Their sizes varied from 0.25µm^2 to 0.85µm^2, falling within typical range of integrin-based adhesion sites. Remarkably, not only did these adhesion sites maintain a consistent orientation, but the internal arrangement of FA and CCACs within them also showed a consistent pattern, pointing existence of polarized mechanical and molecular cues Figure 2a, middle panel).

Through immunostaining experiments, we discerned that certain integrin αVβ5 adhesion sites were distinctly occupied by either FAs or CCACs, while some sites indeed occupied by both complexes Figure 2d). This underscores that these adjacently suited entities inhabit the same adhesion sites. Intriguingly, while the majority of these adjacent CCACs and FAs remained stable over time without evident spatial overlap or interchange (Figure S3d), we observed a marked shift at specific adhesion sites Figure 2a, zoomed in subset in the middle panel). Here, CCACs appeared to exert force to gradually push against and eventually supplanting the FAs. This forceful dynamic suggests a competitive antagonism between CCACs and FAs. Our observations lead us to theorize that, upon extended exposure to Cdk4/6 inhibitors, CCACs might dominate and eventually replace FAs in a manner reminiscent of a relentless tug-of-war.

Building on this, with longer treatment (48 hours), we indeed observed occupation of most adhesion sites by CCACs, and FA expression was almost eliminated Figure 2a, right panel). The CCACs appeared in extended linear arrays and oriented similarly to FA stripes in polarized cells, indicating CCACs might have replaced FAs on integrin αVβ5 adhesion sites over time. However, this duration of treatment caused significant cell death (Figure S3e), possibly indicating the intricate balance between adhesion complexes is essential for cell survival.

To further validate the impact of Cdk4/6 activity on adhesion mode switch and rule out potential side effects of the Cdk4/6 inhibitor, we knocked down Cyclin D1, which is required for Cdk4/6 activation (Figure S3i). Consistent with previous findings, shRNA-mediated downregulation of Cyclin D1 expression also promoted CCAC formation and FA dissociation (Figure S3f,g).

Both chemical and genetic perturbation of Cyclin D1-Cdk4/6 activation resulted in cell cycle arrest at the G1 phase (Figure S3h), raising the question of whether Cyclin D1-Cdk4/6 might be regulating the switch between adhesion modes through impeding the cell cycle progression. The absence of active Cyclin D1-Cdk4/6 results in cell cycle arrest at the G1/S checkpoint, ultimately triggering an exit from the cell cycle into a quiescent state (G0) (102,103). To investigate this, we examined the regulation of adhesion complexes during G0 state by serum-starving cells for 72 hours. Similar to depletion of Cyclin D1-Cdk4/6 activity, we observed a significant increase in CCAC density as cells exited the cell cycle, whereas FAs almost completely dissociated. Reintroducing serum stimulated cell cycle reentry and partially restored the adhesion complex ratio, promoting FA formation and dissociating CCACs cells (Figure S3j). However, this restoration was not to the extent in asynchronized cells. This indicates that, although cell cycle progression was reestablished with introduction of serum (Figure S3h), active Cdk4 is required for complete recovery of adhesion mode. These observations suggest that Cyclin D1-Cdk4/6 may regulate adhesion mode switching not only directly through its kinase effect but also indirectly by influencing the cell cycle progression.

### 3. Membrane tension is the major regulator of adhesion mode switch

Membrane tension has been identified as a key regulator of CCAC formation. Therefore, we hypothesized that alterations in CCACs during the cell cycle may be governed by changes in plasma membrane tension. In support of this hypothesis, our optical tweezer-based measurements of tether force revealed cell cycle-dependent fluctuations in membrane tension force (Figure 3a). The tether force correlates with the energy necessary for the invagination of a clathrin-coated vesicle, elucidating the oscillation we observed in the CCAC to clathrin pit ratio across the cell cycle (Figure 1c). In addition, the inhibition of Cyclin D1-Cdk4/6 activity also resulted in a significant increase in membrane tension (Figure 3b).

**Figure 3:**
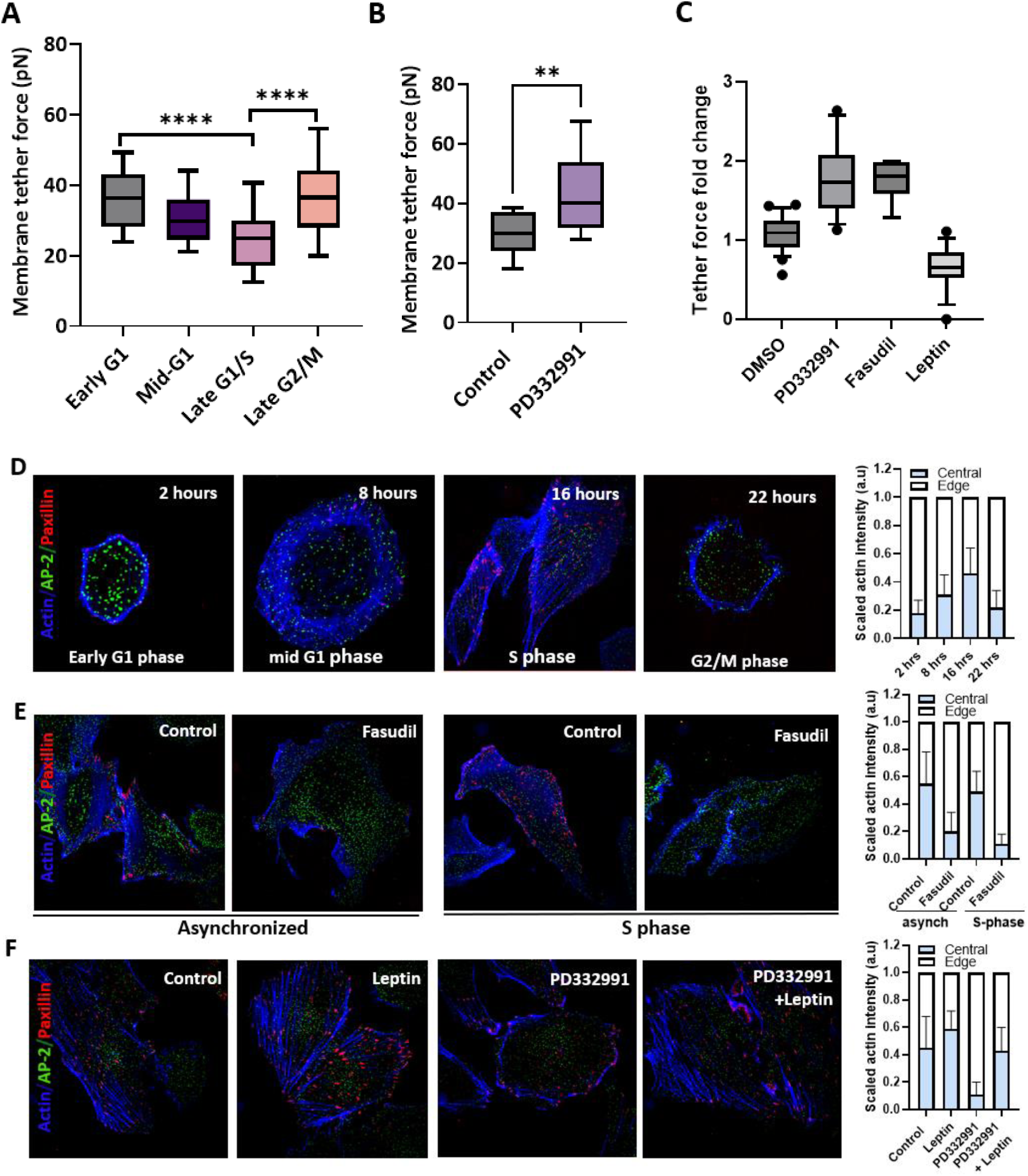
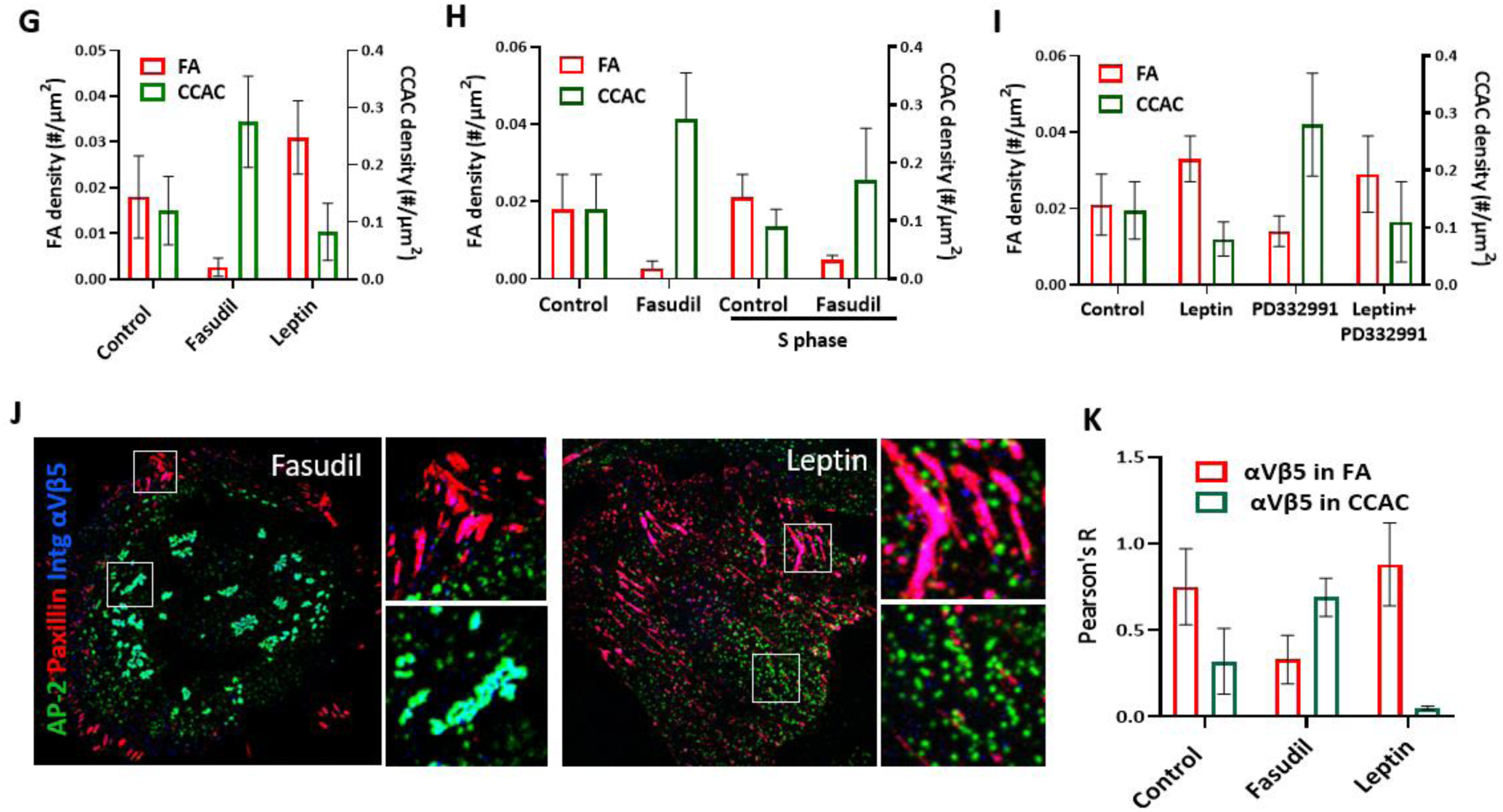
Modulation of membrane tension determines the adhesion mode of cells. **A**) Bar plot illustrating the fluctuation of membrane tether force throughout the cell cycle. **B**) Bar plot displaying changes in membrane tether force in untreated (control) versus PD332991-treated cells. **C**) Bar graph showing the fold change in membrane tether force between untreated (control) cells and those treated with PD332991, Fasudil, or Leptin. Tether force results for Fasudil and Leptin treatments are referenced from Kural et al., 2022. **D**) Phalloidin labeling of cells, genetically edited for AP-2 GFP and stably expressing mCardinal Paxillin, from selected cell cycle stages. The right bar plot presents the normalized distribution of the actin network from the cell periphery to the center. **E**) Phalloidin staining of synchronized cells released into the S phase, both untreated (control) and treated with Fasudil. The right bar plot indicates the normalized distribution of the actin network from periphery to center. **F**) Phalloidin staining of cells untreated (control) or treated with Leptin, PD332991, or a combination of both. The right bar plot depicts the normalized actin network distribution. **G**) Bar plot comparing the densities of FAs and CCACs in untreated (control) cells versus those treated with Leptin and Fasudil. **H**) Bar plot showing the densities of FAs and CCACs in cells from experiment E. **I**) Bar plot illustrating the densities of FAs and CCACs in cells from experiment F. **J**) Immunostaining of AP-2 GFP and mCardinal Paxillin expressing cells with integrin αVβ5, treated with Fasudil or Leptin. The right panels are zoomed-in insets from the boxes in the left images. **K**) Quantification of colocalization between integrin αVβ5 and either CCACs or FAs. All data are represented as means from three or more independent experiments.

Mirroring the mechanoregulatory influence of membrane tension on CCACs, FAs are similarly governed by fluctuations of actomyosin contractility throughout the cell cycle. Similar to many morphogenic events, the cell cycle is punctuated by a periodic transition between the contractility modes, driven by the intracellular reorganization of actin network between cortical and stress fibers in N2A and CHO cells (104). We showed that during G1 to S phase transition, actins reorganize from mostly cortical arrangement into stress fibers and subsequently returns to cortical structure towards the end of G2 phase in SUM159 breast cancer cells (Figure 3d). This observation accounts for the elevated membrane tension observed during early G1 and late G2 phases, as the cortical actin network is known to contribute to membrane tension(105,106). However, during late G1 and S phase, low membrane tension is associated with appearance and consolidation of stress fibers, thereby promoting FA formation. In addition to this, the disassembly of the cortical actin arrangement during G1 renders actins readily accessible, presumably facilitating their role in CCS invagination (Figure 1c, (107), therefore promoting CCAC dissociation.

To further corroborate our hypothesis and establish a link between membrane tension and adhesion mode switch, we utilized both chemical and mechanical approaches to ectopically modulate membrane tension. We recently reported that, fasudil, a Rho kinase inhibitor, increases membrane tension and cortical actins. Conversely, leptin, which increases stress fibers-based actomyosin contractility, reduces membrane tension (Figure 3c) (108). We further verified these results by comparing the phosphorylated ERM (Ezrin, radixin, moesin) proteins in these cells. pERM mediates membrane-to-cortex attachment, therefore, are associated with increased membrane tension (109–112). Consistent with our prior findings, fasudil treatment resulted in a marked increase in pERM localization on the plasma membrane. In contrast, leptin treated cells exhibited primarily cytoplasmic pERM, with no detectable pERM accumulation on the membrane (Figure S4a).

As anticipated, we observed a remarkable switch in densities of the adhesion complexes in response to changes in these forces. Specifically, a fasudil induced elevation in membrane tension promoted CCAC formation, whereas a leptin induced reduction in membrane tension promoted FA formation (Figure 3g). In addition to this, with the elevated membrane tension induced by fasudil, we saw a predominant localization of CCAC to integrin αVβ5. On the other hand, leptin treatment facilitated occupation of αVβ5 sites primarily with FA complexes (Figure 3j, k).

We next assessed the effect of changes in membrane tension on cell cycle regulation of adhesion complexes. We treated cells synchronized to S phase with fasudil and found out that fasudil treatment reversed the adhesion complex ratio and stimulated actin reorganization from stress fibers into cortical actin network (Figure 3e, h). Similarly, in cells arrested at G1 phase due to Cdk4/6 inhibition, leptin treatment was sufficient to reverse the adhesion complex density and stimulated transition from cortical arrangement into stress fibers (Figure 3f, i). We also observed that although fasudil and leptin induced minor shifts in the cell cycle profile, these changes were not significant, indicating that their effects are predominantly exerted through direct modulation of cell membrane mechanics (Figure S4h).

Although leptin stimulates actin stress fiber formation, it doesn’t directly govern actin rearrangement(113). To rule out any indirect effects influencing adhesion complex dynamics during leptin treatment, we exposed cells to Rho A activator I, a recognized promoter of stress fiber formation and its corresponding actomyosin contractility. Similar to leptin treatment, Rho activator I treatment had a counteractive impact on the adhesion complexes: CCACs were reduced by ∼30%, while FAs rose significantly (Figure S4b,e). However, unlike fasudil and leptin, we found that Rho A activator I had significant cytotoxic effects (Figure S4g). Moreover, changes in ratios of the cortical actin to the stress fibers and the subcellular distribution of pERM in Rho activator I treated cells was not as profound as in leptin treatment. This suggests that leptin is a more effective and safer agent for inducing actin rearrangement and consequently, mediating the switch in adhesion mode.

Although we have explored the roles of both actomyosin contractility and membrane tension, it remains unclear which of these mechanical forces is the upstream regulator of the interchangeable deployment of adhesion complexes. To demystify this, our subsequent approach was to directly modulate membrane tension. First, we mechanically induced changes in membrane tension by subjecting cells to a hypoosmotic solution, which increases in-plane tension on the membrane. This intervention led to an increase in CCAC density, a decline in FA density and interestingly reorganization of actin into cortical actin arrangement (Figure S4c, e). Concurrently, we observed a pronounced localization of pERM to the membrane during hypoosmotic shock (Figure S4 f). Our observations resonate with prior studies highlighting the reciprocal regulation between cortical contractility and membrane tension(87,105,114–116). In conjunction with these studies, our data indicate that while cortical tension contributes to increased membrane tension, a rise in in-plane membrane tension also actively promotes cortical actin organization.

As an alternative approach to modulate membrane tension directly, we treated cells with methyl-beta-cyclodextrin (MBCD), a compound known for depleting cholesterol and increasing cell membrane tension. This treatment mirrored our earlier findings, revealing a surge in CCAC density and a drop in FAs (Figure S4 d,e). Further, MBCD also induced cortical actin formation and pERM clustering on the membrane (Figure S4 f). Interestingly, upon removal of MBCD and its replacement with normal media, as well as post-restoration of tension following hypoosmotic shock, we detected reformation of FAs accompanied by rearrangement of stress fibers (Figure S4c). These data collectively indicate that change in membrane tension serves as the primary driving force behind the adhesion mode transition, with the actin arrangement either synergizing with or counteracting this mechanism.

### 4. Membrane tension regulates cell spreading and migration via coordinating adhesion complexes

Coordination of membrane tension and actomyosin contractility is essential for several processes including cell spreading and migration(18,88,117–119). Notably, cancer cells, characterized by enhanced proliferation, spreading, and migration, tend to have softer membranes compared to non-cancerous cells (Ref). Based on this observation, we hypothesized that the mechanical state of a cancer cell, particularly its adhesive behaviors, plays a central role in its increased capacity for spreading and migration.

To test this, we first compared the spread area and migration speed of SUM159 cells during the cell cycle. We found that cell cycle stages associated with increased membrane tension —and consequently, a predominance of CCACs at adhesion sites—correlated with reduced spreading and migration abilities (Figure 4a-c). In contrast, during the late G1 and S phases where membrane tension is low, stress fibers-based contractility is high and therefore adhesion sites are dominated by FAs, we observed significant increase in cell spreading and migration. Interestingly during the early G1 and G2 phases-the cell cycle stages which corresponds to switch in adhesion modes-we noted a discernible inflection point in the trajectory of cell migration speed, further indicating the counteractive roles of CCACs and FAs in cell migration (Figure 4a, S5a, pointed by arrows). Reinforcing these findings, pharmacological inhibition of Cdk4/6 activity and cell cycle exit to quiescent (G0) state, which increase membrane tension, substantially reduced the migration abilities of a cell (Figure S5 b).

**Figure 4:**
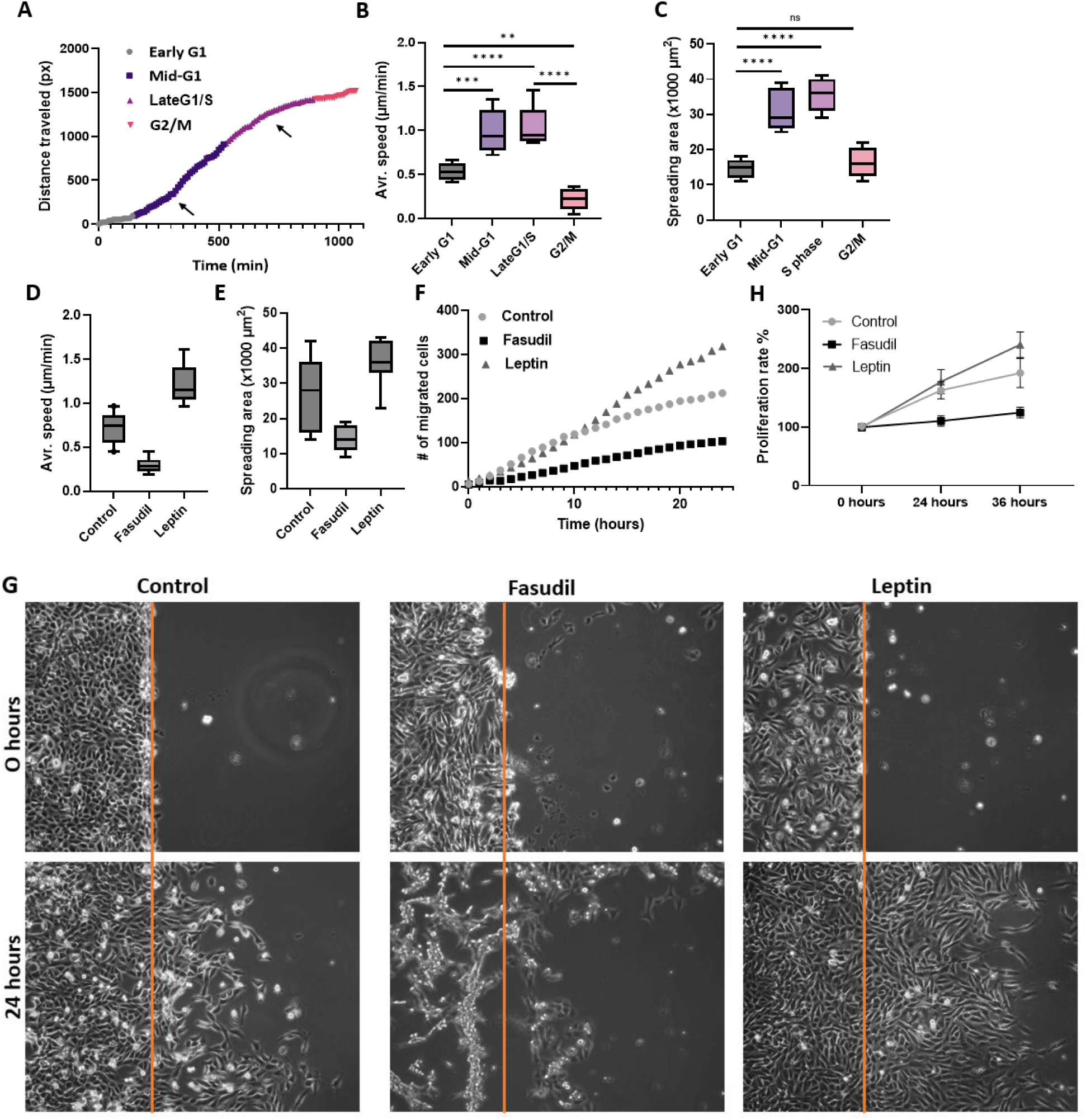
Membrane tension regulates migration and spreading of cells. **A**) Linear plot depicting the distance traveled by cells over time throughout the cell cycle, with arrows highlighting the inflection points. **B-C**) Bar plots showing changes in average speed and spread area of cells at specified time points. **D-E**) Bar plots comparing the average speed and spread area of cells either untreated (control) or treated with Fasudil or Leptin. **F**) Bar plot indicating the number of cells that migrated into a ‘wound’ area, comparing untreated cells with those treated with Fasudil or Leptin, as shown in figure G. **H**) Linear plot illustrating the proliferation rates of control cells versus those treated with Fasudil or Leptin. All data are presented as means from three or more independent experiments.

We observed similar results when membrane tension and actomyosin contractility were modulated externally. Fasudil treatment reduced spreading and migratory capabilities, whereas leptin expanded the cell spread area and enhanced motility (Figure 4d-g). Furthermore, we observed significantly more blebbing and protrusions in leptin treated cells (Figure 4g, right bottom panel). Collectively, these observations emphasize the role of the intricate interplay between membrane tension and actomyosin contractility in determining a cell’s adhesive modalities, which subsequently influence its spreading and migration dynamics.

In addition to the mode of adhesion, the strength of adhesion plays critical role in cell spreading and migration(120). It is known that strong and stable adhesion prevents the cell from releasing its cytoskeleton-ECM linkages, whereas weak and transient adhesion does not generate the contractile force necessary for cell movement(121–125), indicating that an intermediate state of adhesion is most favorable for efficient migration and spreading. To delve deeper into understanding adhesion complexes’ dynamics, we treated cells with cilengitide. This compound acts as competitive ligand mimetic inhibitor to integrins(126) and does not disrupt existing integrin binding sites. Instead, it selectively impedes the formation of new integrin interactions, thereby can be used to assess the stability of adhesion complexes. Live imaging of cilengitide treated cells showed that FAs persisted longer than CCACs (Figure S5 c), indicating FAs are more stable structures. The transient or weak interaction of CCACs with the ECM is presumably insufficient for establishing robust anchoring points necessary for effective membrane protrusions, which explains reduced cell spreading and migration. On the contrary, cells treated with a high concentration of leptin (100nM) and Rho activator I (10µl/ml), also resulted in stalled migration (Figure S5d), presumably due to highly stabilized FAs and stress fibers. This indicates that cells require a fine-tuned balance between CCACs and FAs to efficiently spread and migrate.

In addition to their role in cell spreading and migration, regulation of cell adhesion is imperative for proper cell cycle progression and proliferation(13,127–134). While short term treatment of cells with fasudil (20µM, 2 hours) and leptin (30nM, 24 hours) did not markedly alter cell cycle profile (Figure S4h), interestingly, extended exposure elicited substantial changes in cell proliferation rate. Time-lapse imaging of cells for 36 hours showed that, Fasudil (1µm) treatment notable decreased proliferation, whereas leptin (30nM) enhanced it (Figure 4g, S5e).

Based on these observations, we speculate that when membrane tension is high, CCACs act as an antagonist of FAs at integrin αVβ5 based adhesion sites. Through their weak and transient interaction with ECM, CCACs act as restrains on cell proliferation, spreading and migration. However, when membrane tension is low, CCACs dissociate from these adhesion sites, allowing the formation of FAs and generation of forces necessary for these cellular behaviors (Figure 5).

**Figure 5:**
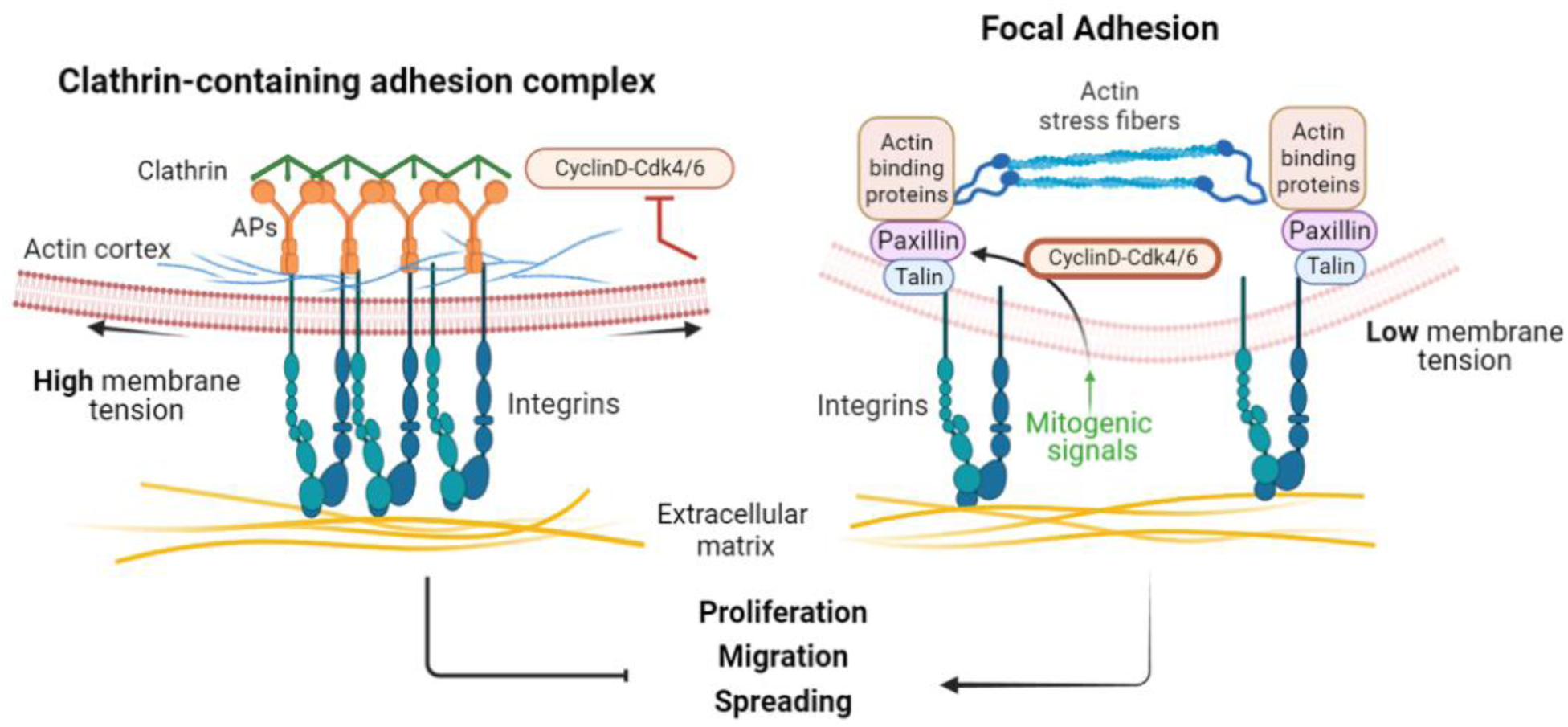
Membrane tension-regulated adhesion mode is essential to determine the extent of proliferation, migration and spreading. **A**) The schematic summary of the report where high membrane tension, along with cortically arranged actins and reduced Cyclin D-Cdk4/6 activity results in occupation of integrin αVβ5 adhesion sites with CCACs. This prevents cells migration, spreading and proliferation. APs: Adaptor proteins **B**) When membrane tension drops, actins organize into stress fibers and in presence of active Cyclin D-Cdk4/6, CCACs dissociate, allowing formation of FAs on integrin αVβ5 adhesion sites. This facilitates migration, spreading and proliferation.

## Discussion

Membrane tension is increasingly appreciated to be a key regulator of many cellular functions(90,112,118,119,135–139). Here, we found that change in membrane tension exerts an upstream control in adhesion modes of cells eventually affecting multiple cellular process which demand extensive membrane remodeling. Our findings indicate that two distinct adhesion complexes-CCACs and FAs-are not only utilized interchangeably but also compete in their deployment. This competitive and interchangeable utilization of the adhesion complexes is coordinated by the intricate interplay of membrane tension and actomyosin contractility, ultimately influencing the extent of cell proliferation, spreading and migration in cancer cells.

Cancer cells exhibit enhanced proliferation and migration(140), This is partly due to their softer plasma membrane and disrupted actin dynamics that allow rapid and extensive morphological changes (141–152). In a multicellular setting, higher cortical contractility in epithelial cells reinforces cell-cell adhesions and contribute to preservation of tissue integrity. Whereas enhanced stress-fiber-based contractility and reduced cortical stiffness in mesenchymal cells, enable them to detach and migrate from the primary tumor into ECM during metastasis(143,153–155). Our data corroborates these concepts and reveals that decreased membrane tension along with increased stress fiber formation enhances the aggressive behaviors observed in cancer cells via promoting occupation of integrin αVβ5 based adhesions sites with FA. However, elevated membrane tension and cortical actin organization facilitates occupation of integrin αVβ5 based adhesions sites with CCAC and reduces malignant features of cancer cells.

Alterations in membrane tension not only dictate adhesion modes but also lead to dysregulated endocytosis in cancer cells. Our findings indicate that reduced membrane tension is associated with an increase in clathrin pit formation (Figure 1c), influencing CME dynamics. The enhancement in CME dynamics facilitates the uptake of nutrients, signaling receptors, and adhesion complexes, furthering cancer progression (156–159). Consequently, targeting endocytosis systematically has been recognized as a promising anticancer strategy (160,161). Beyond molecular interventions in CME, manipulating the mechanical properties of cancer cell membranes has shown potential in reducing malignancy (152,162–164). For instance, it was shown that increasing membrane tension prevented endocytosis of death receptors, thereby enhancing sensitivity of cancer cells to immune cell-mediated destruction (108,165). Therefore, increasing membrane tension might not only sensitize cancer cells to apoptosis but also inhibit clathrin pit and FA formation, further impeding cancer progression.

Moreover, consistent with others, we observed an inverse relationship between the strength of membrane-to-cortex attachment and cancer malignancy(166–169). We found that pERMs are translocated to plasma membrane when cells were treated with fasudil, MBCD or exposed to hypoosmotic shock (Figures S5 a,f), indicating positive feedback loop between membrane tension and localization of pERM to the membrane. High levels of pERM on cell surface are reported to prevent BAR/F-BAR proteins to deform the plasma membrane. Constitutively active ezrin promotes pERM translocation to the plasma membrane, reducing metastasis and proliferation (170,171), while dephosphorylated ERM detaches from the membrane and facilitates lamellipodia formation and cell migration in breast cancer cells (172). Additionally, functional perturbation of ERM proteins impedes CCP maturation and reduces rate of endocytosis(173). Hence, these observations, along with our own, imply that the translocation of pERM to the membrane promotes the formation of CCACs rather than clathrin pits by hindering membrane curvature. This could occur either by directly obstructing curvature-generating proteins or by increasing membrane tension through a reinforced connection to the cortical actin.

In addition to mechanoregulations of adhesion complexes, we identified CyclinD1-CDK4/6 – an oncogene notably overexpressed in most tumor cells – as an essential upstream regulator influencing adhesion mode of the cell. We found that inhibition of CyclinD1-CDK4/6 prevented CCAC to FA switch and induced membrane tension elevation presumably via triggering G1/S checkpoint Figure 2a,b and 3b). This observation is consistent with previous studies showing that the failure to degrade adhesion complexes in a timely manner or alterations in membrane tension due to high osmolarity or reorganization of actin network can induce cell cycle arrests (13,174,175). While further investigation is necessary to determine the reciprocal relationship between checkpoint activation and tension induced adhesion mode switch, our data further underscore the rationale of targeting Cdk4/6 activity as a therapeutic strategy in oncology (176,177).

The observed inverse crosstalk between CCACs and FAs which is mediated by the competitive redistribution of αVβ5 integrins also exists between other adhesion complexes (72,74,178–180). For instance, competitive interactions of hemidesmosomes and FAs with integrin α6β4 determines the invasive phenotype in cancer cells, where presence of integrin α6β4 in hemidesmosomes is correlated with a diminished invasive capability(181–184). Moreover, hemidesmosomes contribute to regulate FA and CCAC stability in keratinocytes via controlling their localization to αVβ5 integrin in tension dependent manner(72,185). Similar inverse dynamics is observed between invadosomes and FAs: invadosome assembly is favored when FA and stress fiber-based actomyosin contractility are low(179,186) (Kedziora 2016). Conversely, the dissolution of invadosomes correlates with the formation of β3-integrin-rich FAs, impairing the ECM remodeling during invasion(187). These observations underscore that, beyond the mere availability of integrins, the affinity of integrins to adhesion complexes is critical in determining the adhesion mode of a cell. Drawing on these findings, we hypothesize that CCACs exhibit a greater affinity for integrin αVβ5 than FAs (Figure S5c), therefore predominate αVβ5 adhesion sites and prevent FA assembly. This hypothesis is substantiated by studies showing that overexpression of β5 facilitates the formation of CCACs, whereas the depletion of β5 impairs CCACs without similarly affecting FAs(Baschieri et al., 2018b; J. G. Lock et al., 2018a), indicating higher affinity of β5 to CCACs.

Adhesion dynamics is also affected by binding strength of adhesion complexes to ECM and cytoskeletal fibers. For efficient spreading and migration, strong yet dynamic anchoring adhesion points are required(188–190). Our data suggest that CCAC mediated adhesions are weaker and more transient (Figure S5c), insufficient to drive forces for cellular protrusions. This is consistent with observations that adhesion strength is at its minimum during mitosis (191), where CCACs are the predominant adhesion complexes, rendering cells immobile (192,193). Furthermore, not only the overall force exerted by the cells, but the traction force normalized by the area are cell cycle regulated. Traction force tend to increase as cells progress through G1, before reaching a plateau in S phase, and then decline during G2(194,195) which mirrors FA expression (Figure 1a,b). This implies that when bound to αVβ5 adhesion sites, FA links the stress fibers to ECM and exert localized and optimal forces conducive to efficient cell spreading and migration. In contrast, αVβ5 integrin sites occupied by CCACs fail to serve as robust anchoring points, as they are not tethered to actomyosin fibers but are influenced solely by the overall mechanical state of the membrane. Although these hypotheses have valid assumptions to build on, further detailed assessments are needed for quantification of CCAC versus FA driven cell to ECM attachment strength.

Collectively, our research demonstrates that elevated membrane tension imparts a dominant negative effect of CCACs on FAs at integrin αVβ5 adhesion sites. This results in weaker and more transient ECM interactions, thereby inhibiting cell proliferation, spreading, and migration. Conversely, when membrane tension is reduced, CCACs release their hold on these adhesion sites, facilitating the formation of FAs and the exertion of mechanical forces essential for cellular protrusions (Figure 5). In summary, our findings provide novel insights into the biomechanical regulation of CCACs and FAs, highlighting their respective roles in either suppressing or promoting cancer cell progression. The data we present serve as a benchmark for future research into the coordination of diverse adhesion complexes across various biological contexts. Furthermore, examining the levels of CCACs and FAs in cancer cells of varying degrees of malignancy, as well as their healthy counterparts could be insightful. Such an investigation may contribute to the identification of novel biomarkers for cancer malignancy. Overall, these studies have the potential to significantly advance our understanding of the complex biological mechanisms that contribute to cellular processes demanding extensive membrane remodeling.

## Acknowledgements

This work was funded by Pelotonia Fellowship awarded to U.D. and E.T.C., NSF Faculty Early Career Development Program (award number: 1751113) and NIH R01GM127526 to C.K. Any opinions, findings, and conclusions expressed in this material are those of the authors and do not necessarily reflect those of the Pelotonia Fellowship Program or the Ohio State University. We also thank Dr. Dmitri Kudryashov, Dr. Elena Kudryashova from the Ohio State University for helping us with the viability assays.

## Author contributions

U.D. oversaw the project, designed, and optimized the experiments and analyzed the data. C.K and U.D. prepared the manuscript. U.D. performed cell cycle synchronization and profiling; adhesion complex analysis with immunostaining, immunoblotting and live imaging experiments; actin organization assessment; pERM and integrin staining experiments; genome editing SUM159 cells for AP-2 Halo expression and shRNA transduction experiments; migration, proliferation, and viability assays. V.A. helped with cell cycle profiling experiments. Y.M. and E.C. performed tether force measurements with C-Trap. E.C. prepared PDMS pieces for wound healing assays. T.W. and C.K. helped with CCAC quantification methods. A.A. helped with CCAC analysis during cell cycle.

