## Supplementary data for "Tension-induced adhesion mode switching: the interplay between focal adhesions and clathrin-containing adhesion complexes"

**Supplementary Figure 1:**

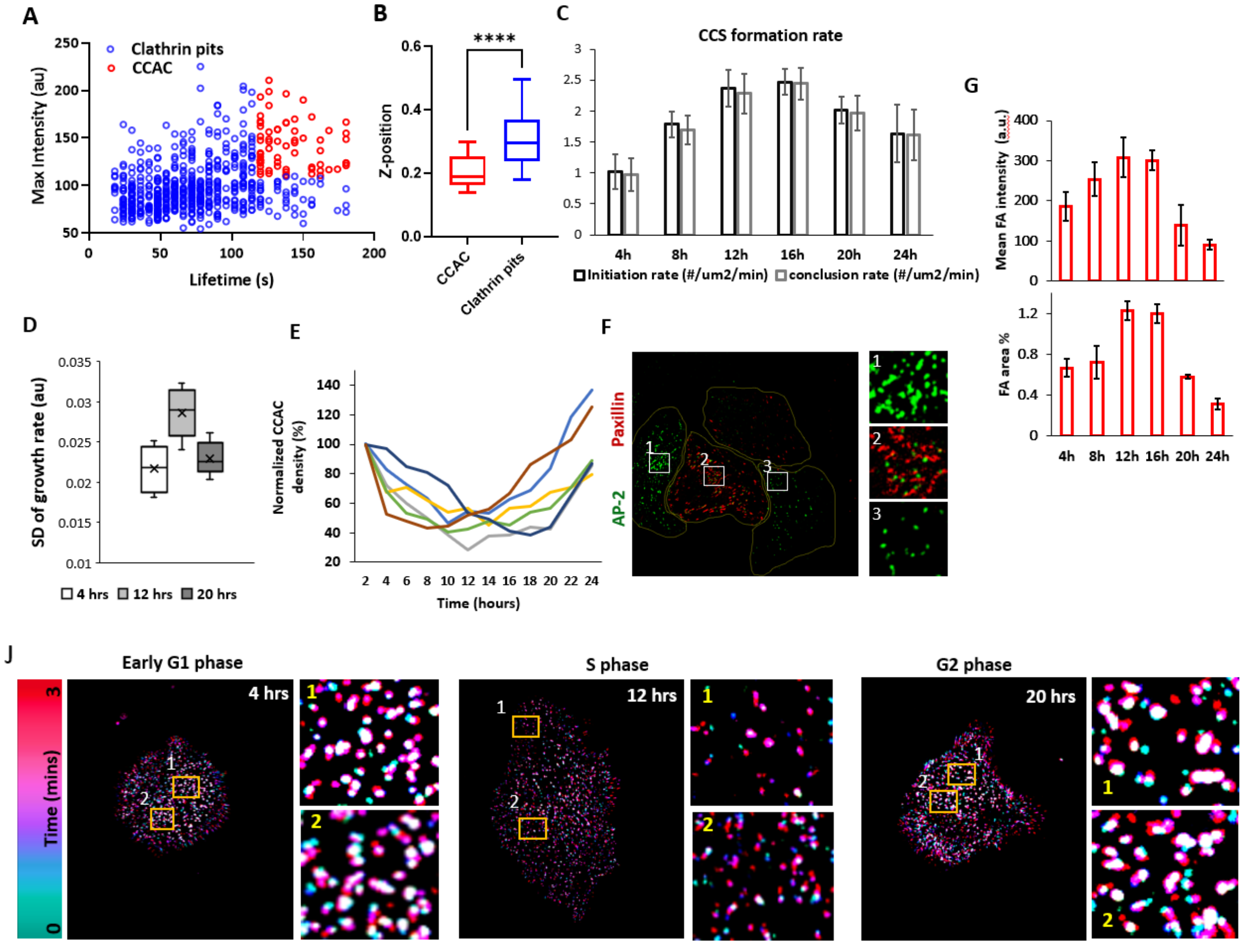

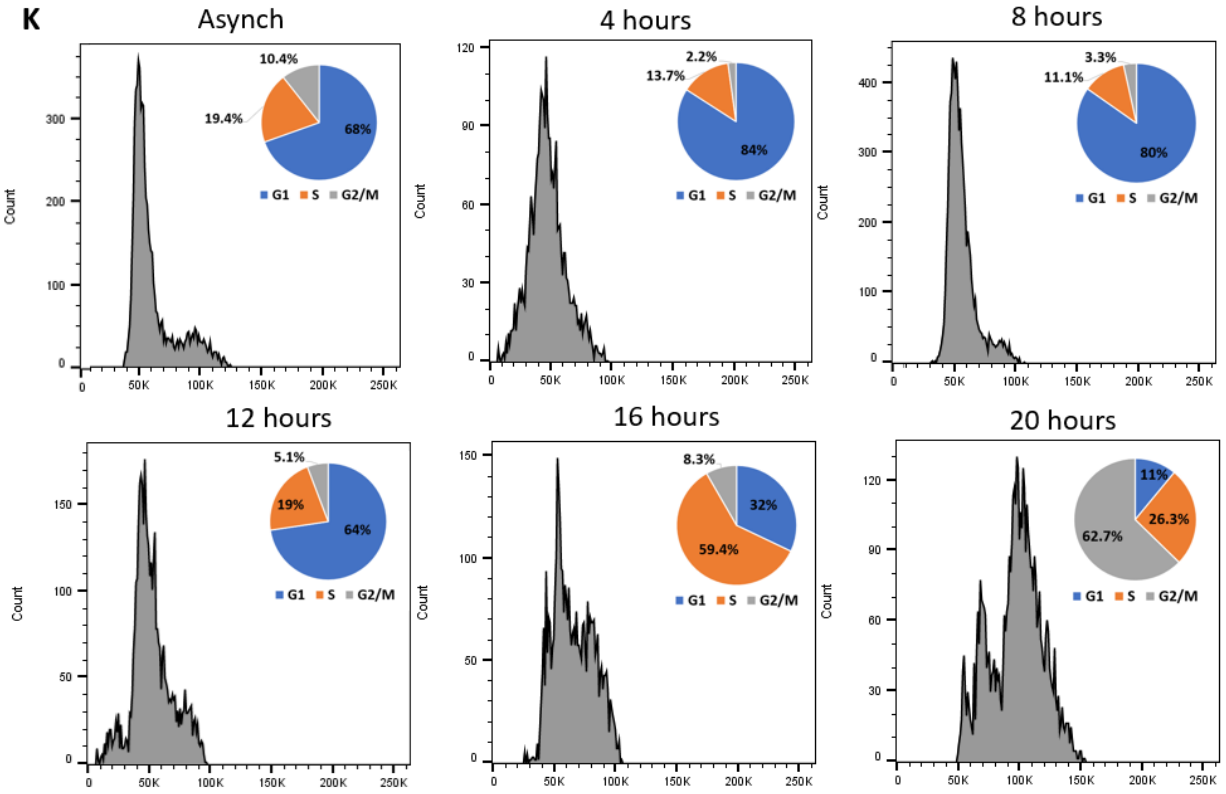


**Supplemental figure 1: Quantification of CCAC and FA abundance:** A) Scatter plot shows the max intensity versus the lifetime of traced CCS with CME analysis from 3 minutes long movies. Dashed red lines are the thresholds applied to a lifetime (>120 seconds) and intensity (>100). The upper right of the plot, CCS structures labeled with red are the CCAC structures. B) The whisker plot shows the s z-position of the CCAC and clathrin pits. C) Immunolabeling of integrin B1, b3 and b5, paxillin and AP-2. D) Quantification of integrin localization to CCAC (thresholded CCS) and FA (all paxillin). E-F) Bar plot showing standard deviation (SD) of growth rate and the mean (±SD) CCSC formation and dissociation rates plotted versus time. G) Mean area and intensity of FA at indicated times points. H) CCAC density of 8 different cells monitored throughout the cell cycle. I) Representative image for relative CCAC and FA expression in asynchronized cells. J) Temporal color code of CCS traced for 3 minutes at indicated time points. The color code shows the lifetime of structures. K) Flow Cytometry of synchronized cells at indicated times points. L) Colocalization of integrin B5 with FA and CCAC complexes at indicated times points of cell cycle. M) Quantification of the colocalization.

**Supplemental figure 2:** CCAC colocalizes with avb5 but not with b1 and b3

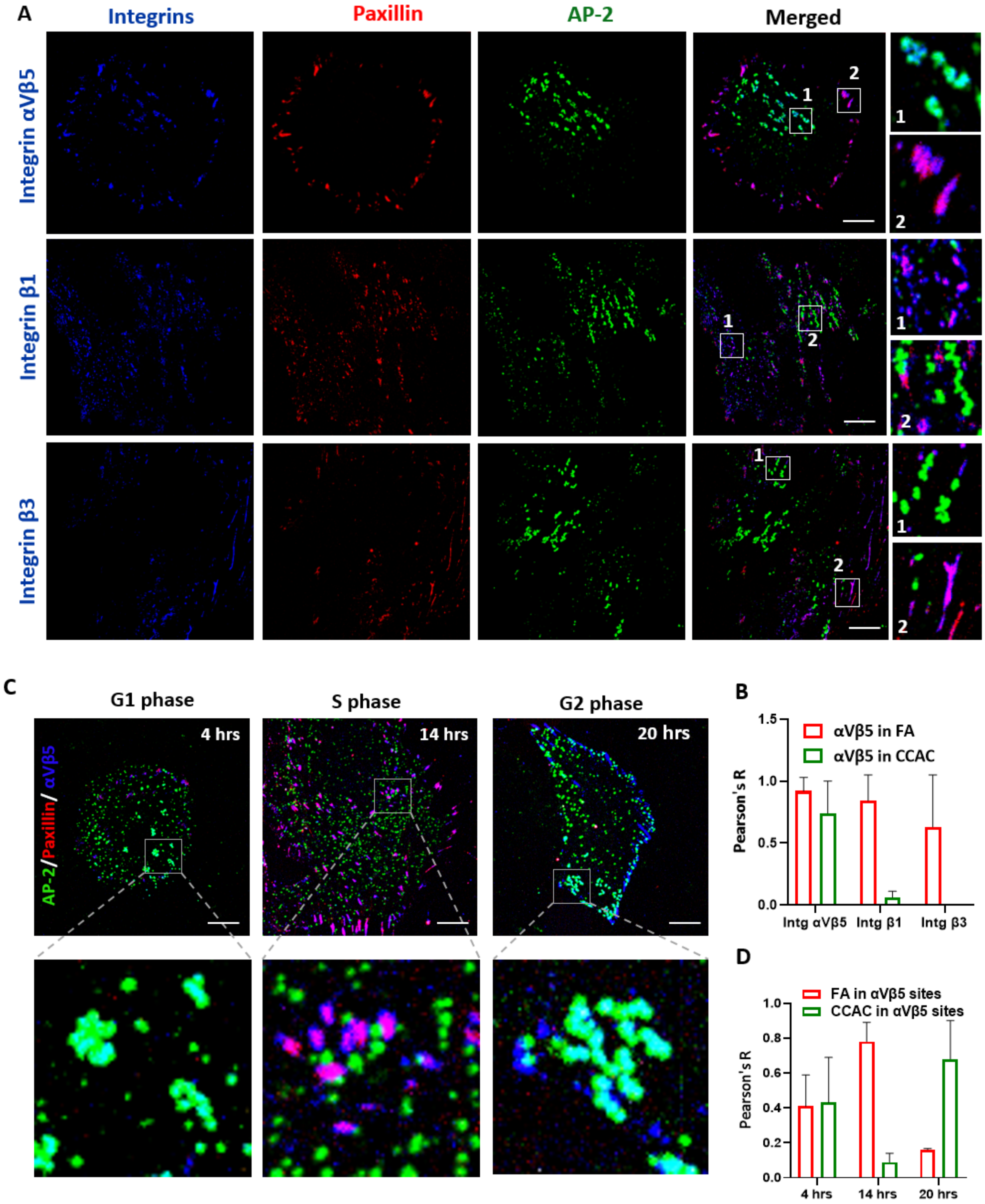

**Supplementary figure 3:**

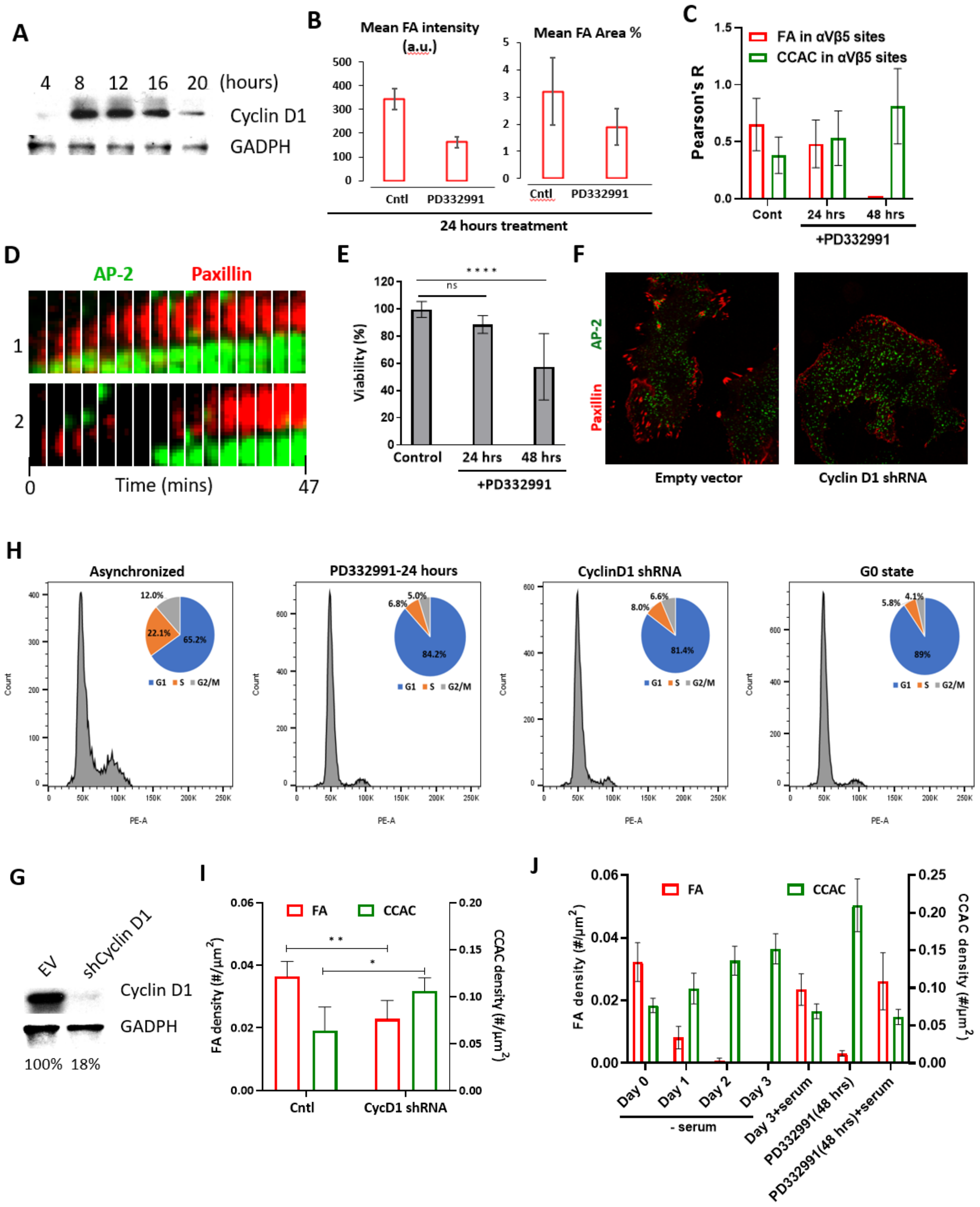

**Supplementary Figure 4:**

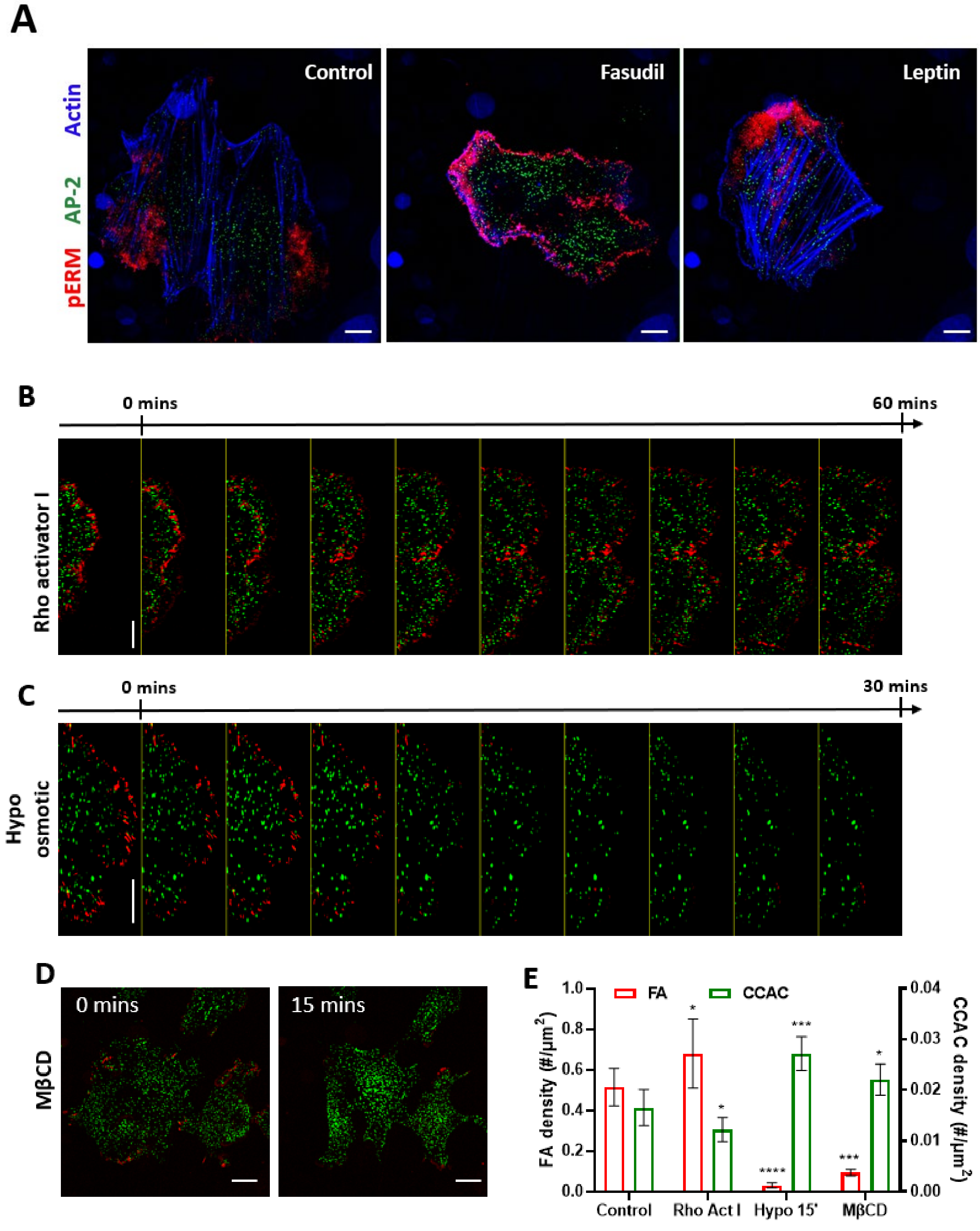

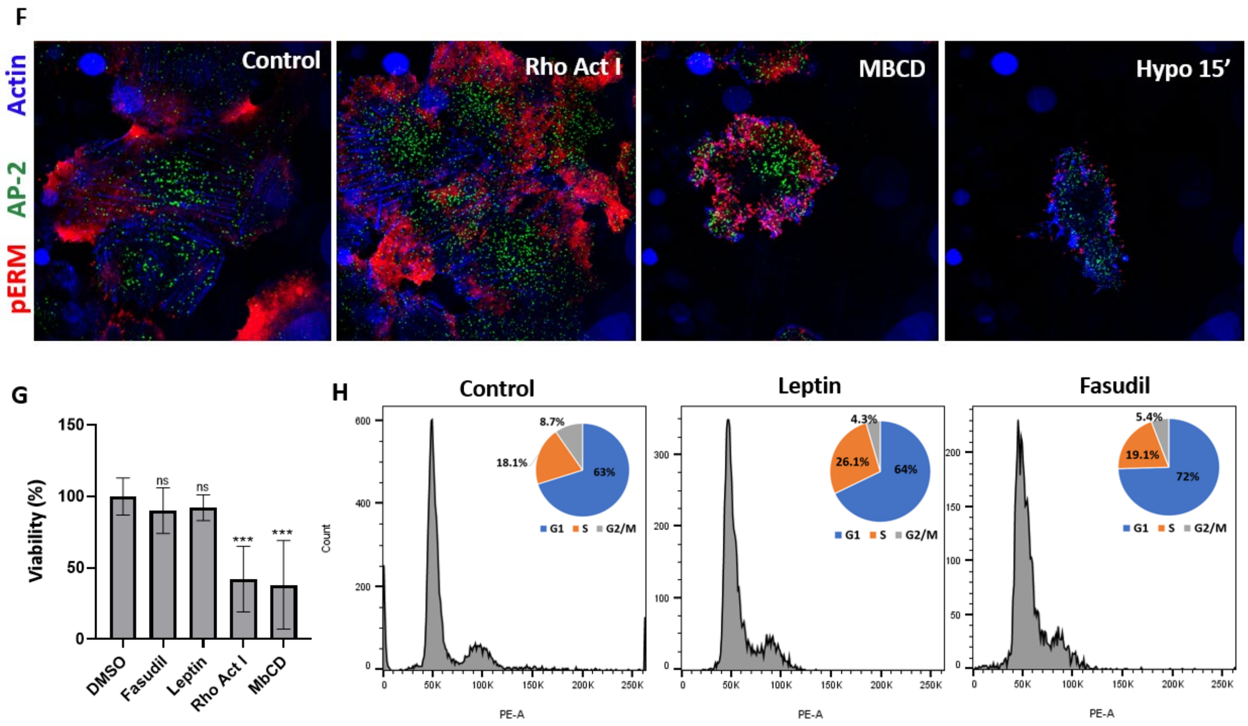
F
Actin
AP-2
pERM
Control
Rho Act I
MBCD
Hypo 15’
G
Viability (%)
150
100
50
0
ns
ns
***
***
DMSO
Fasudil
Leptin
Rho Act I
MbCD
H
Control
Leptin
Fasudil
Count
PE-A
8.7%
18.1%
63%
4.3%
26.1%
64%
5.4%
19.1%
72%
G1
S
G2/M
0
50K
100K
150K
200K
250K

**Supplementary Figure 5:**

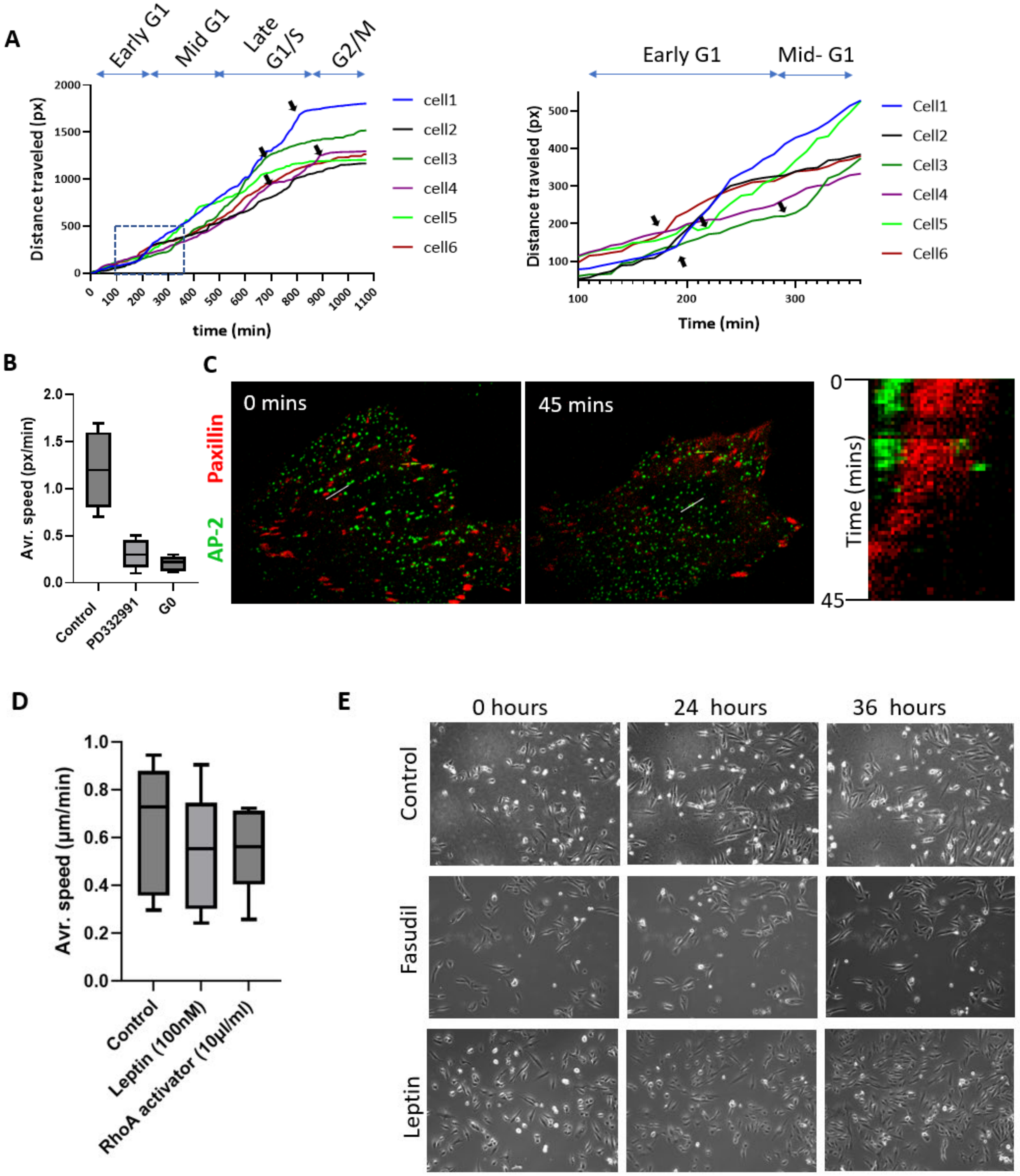
